# Effect of ethanol exposure *in utero* on infant mice gut microbiotas assessed by nanopore and illumina sequencing

**DOI:** 10.1101/2022.12.09.519727

**Authors:** Cristiano Pedroso-Roussado, Fergus Guppy, Nigel Brissett, Lucas Bowler, Joao Inacio

**Author notes:** Address correspondence to Cristiano Pedroso-Roussado,.

## Abstract

The gut microbiome plays a vital role in host homeostasis and understanding of its biology is essential for a better comprehension of the etiology of disorders such as Foetal Alcohol Spectrum Disorder. Foetal Alcohol Spectrum Disorder represents a cluster of abnormalities including growth deficiencies and neurological impairments, which are not easily diagnosed nor treated. Here the effect of ethanol exposure *in utero* on the gut microbial profiles of 16 infant mice (nine exposed *in utero* and seven non-exposed) was assessed by targeted nanopore sequencing and Illumina sequencing approaches. The nanopore sequencing was implemented using MinION system targeting PCR-amplified amplicons made from the full-length 16S rRNA gene. The Illumina sequencing was performed using Miseq system targeting the V3-V4 region of the 16S rRNA gene. Ethanol exposure did not affect the microbial profiles. Several low prevalent taxa, like *Akkermansia muciniphila*, were detected but further studies must be performed to detail the effect of ethanol exposure to these taxa since no clear pattern was detected throughout this study.

**Importance:** Detailed knowledge about the interactions between gut microbes and the developing nervous system is still scarce. Foetal Alcohol Spectrum Disorder represents a clinically relevant set of conditions with cumbersome diagnostic and treatment. In this work the microbial profiles of infant mice gut exposed to ethanol *in utero* were analysed through third-generation Illumina and optimized next-generation nanopore sequencing technologies. The fungal (albeit not detected) and bacterial microbial profiles here obtained through nanopore and Illumina sequencing represent a technological and biological advancement towards a better comprehension of the microbial landscape in Foetal Alcohol Spectrum Disorder at early post-natal periods.

## Introduction

According to the World Health Organization, 283 million people worldwide had a diagnose related with alcohol use disorders in 2018 (1). Firstly, ethanol consumption causes dysbiosis in gastrointestinal tract with observed increase of Gram-negative bacteria (2) (3), and decrease of short-chain fatty acids (SCFA)-producing bacteria (4). Secondly, endotoxins produced by Gram-negative bacteria compromise intestinal barrier integrity causing higher permeability (5) (6). Enhanced permeability allows bacterial cells and metabolites to cross and enter the portal and the systemic circulation system, causing damage elsewhere in the body (7). In humans, the effect of ethanol has consequences on the gut landscape and on the microbial composition (4) (8) (9) (10) (11) (12), and it also has a prejudicial effect on the brain, and cause liver lesions by the disruption of the intestinal barrier (13). However, there are few clinical studies about the impact of gut microbiota in ethanol-dependent humans (14). At early age, infants acquire their microbiome most probably from their mothers during pregnancy, then it develops until relative maturation in the first three years of life (10). Prenatal environment and early postnatal colonization are crucial for a normal general and organ-specific development (15). However, despite the increasing amount of data, there is still no strong evidence about the potential influence of the gut microbiota in alcohol use disorder and neonatal immune, physiological, and neurological development (16). The gut microbiome plays a vital role in host homeostasis and understanding of its biology is essential for a better comprehension of the etiology of disorders such as Foetal Alcohol Spectrum Disorders (FASD). FASD represent a cluster of abnormalities including growth deficiencies and neurological impairments, which are not easily diagnosed nor treated (17) (18). Overall, the effects of prenatal alcohol exposure on the neurodevelopment can be observed in a myriad of neurological processes, for instance involving microglia and astrocytes, promoting above normal levels of inflammatory response during the central nervous system development (19). Third-generation technologies such as nanopore sequencing whilst showing considerable promise still need to be further developed to improve the reliability of their sequencing outputs. So far, scientists have been employing a two-way sequencing approach to validate and/or complement nanopore sequencing studies together with more mature sequencing technologies, such as short-read Illumina sequencing. However, there are several technical challenges to be addressed in nanopore sequencing approaches, mostly related to the non-standardization of experimental and bioinformatics protocols. Indeed, our knowledge of the composition and function of the whole gut microbiota lacks a comprehensive description, with particular significance in a clinical context where appropriate diagnostics and therapeutics may be lacking, such as FASD. Since there is no consensus regarding foetal and neonatal gut microbiota transmission and colonization, the impact of stressors like alcohol consumption occurring during pregnancy needs an urgent analysis. Recent advances in sequencing technologies have emphasized questions about the microbial profiles obtained when different approaches are tested. Such questions are mainly due to the genomic target used for sequencing and differences in the bioinformatic pipeline used (20) (21) (22) (23) (24). Here the effect of ethanol exposure *in utero* on the gut microbial profiles of 16 infant mice (nine exposed in utero and seven non-exposed) was assessed by targeted nanopore sequencing and Illumina sequencing approaches. The nanopore sequencing was implemented using MinION system targeting PCR-amplified amplicons made from the full-length 16S rRNA gene. The Illumina sequencing was performed using Miseq system targeting the V3-V4 region of the 16S rRNA gene.

## Results

### Nanopore and Illumina sequencing quality overview

PCR was performed to generate fungal ITS amplicons from the infant mice samples. However, no amplification was detected by gel electrophoresis from any sample (data not shown). This observation suggests that fungal members were absent or in very low abundance in the infant mice gut. Following the apparent absence of a gut mycobiome in our sampled infant mice, the remaining work focused on the bacterial gut microbiota of the same infant mice. In order to gather the highest number of sequenced reads possible, four independent nanopore sequencing runs were performed (Table 1).

**Table 1.** Nanopore sequencing performance parameters of 4 independent sequencing runs of infant mice faecal samples exposed to ethanol in utero and control samples.

| Barcoded samples ID<sup>1</sup> | No. reads | No. bases (bp) | Mean read length (bp) | Read length N50 (bp) | Mean read quality (Qscore) | Percentage of reads above quality cutoff (%) |  |
| --- | --- | --- | --- | --- | --- | --- | --- |
|  |  |  |  |  |  | Q10 | Q12 |
| Run A | 42,073 | > 67 million | 1,606 | 1,616 | 10.3 | 63.4 | 3.7 |
| Run B | 625,975 | > 1,0 billion | 1,615 | 1,621 | 10.4 | 62.2 | 9.7 |
| Run C | 557,179 | > 899 million | 1,614 | 1,623 | 10.1 | 52.7 | 3.9 |
| Run D | 989,109 | > 1,5 billion | 1,603 | 1,614 | 10.5 | 67.9 | 9.6 |

<sup>1</sup> – In Run A, two control samples were sequenced; In Run B, three control samples were sequenced; In Run C, four test and two control samples (one of which was a repeated sample) were sequenced; In Run D, five test and 1 control samples were sequenced.

In respect to Illumina sequencing, a total of 1,988,055 reads were obtained, with a total yield of 1,132,178 kilobases. All reads passed the default Illumina chastity filter procedure and more than 60% of the reads had passed Q30 and the mean Q obtained was 29.2. As expected, Illumina sequencing retrieved a decreased average error-rate in comparison with nanopore sequencing (~0.01% *vs* ~9%).

### Comparison between the microbial profiles of infant mice guts exposed to ethanol *in utero*

*Firmicutes* was the most represented phylum (53%), followed by *Bacteroidetes* (37%), *Verrucomicrobia* (10%), and *Deferribacteres*, which did not exceed 1% of the assigned reads in the ethanol-exposed group analysed by the targeted nanopore sequencing approach. At the genus level, the most prevalent genera were *Lactobacillus* (19%), followed by *Duncaniella* (17%), *Muribaculum* (16%), *Limosilactobacillus* (13%), *Faecalibaculum* (13%), and *Akkermansia* (10%), and the remaining genera had less than 3% of relative abundance. A total of 21 species were assigned. The most represented species were *Duncaniella muris* (17%), *Lactobacillus johnsonii* (17%), *Muribaculum intestinale* (16%). *Limosilactobacillus reuteri* (13%), *Faecalibaculum rodentium* (13%), and *Akkermansia muciniphila* (10%) and the remaining 14 species had less than 3% of percentual relative abundance.

In the control group, *Firmicutes* was the most represented phylum (46%), followed by *Bacteroidetes* (44%), *Verrucomicrobia* (10%), and *Deferribacteres*, that accounted for less than 1% of the reads. At the genus level, the most represented genera were *Muribaculum* (20%), followed by *Duncaniella* (20%), *Lactobacillus* (16%), *Akkermansia* (10%), *Limosilactobacillus* (9%), *Faecalibaculum* (5%), and *Lachnoclostridium* (5%). The remaining genera had less than 3% of relative abundance. In the control group, a total of 27 species was observed. The most represented was *M. intestinale* (20%), followed by *D. muris* (20%), *L. johnsonii* (14%), *A. muciniphila* (10%), *L. reuteri* (9%), and *F. rodentium* (5%). The remaining 21 species had less than 3% of percentual relative abundance. These microbial profiles here observed does not completely match other reports (10) (13) (25) (26). However, the changes are not easily comparable due to the different experimental conditions performed, such as mice model used, diet, and ethanol administration regimen (28). Specifically, neither *Prevotella, Pediococcus, Faecalibacterium, Bifidobacterium, Escherichia*, nor *Staphylococcus* were detected in this study, however they were detected in another study (25). Nanopore and Illumina sequencing approaches retrieved different microbial profiles from the same infant mice gut faecal samples collected from mice exposed and non-exposed to ethanol *in utero* (Figure 1).

**Figure 1.**
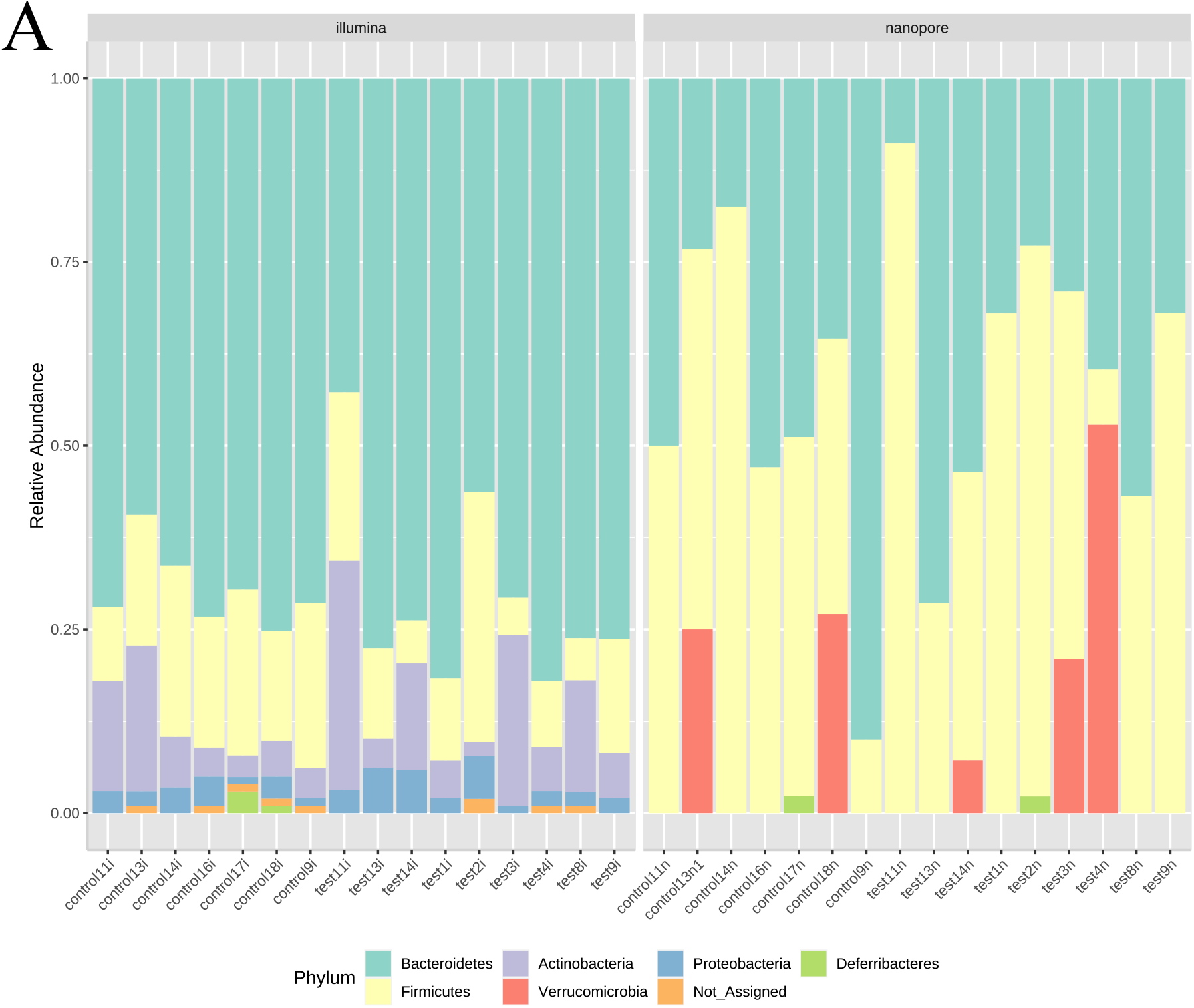
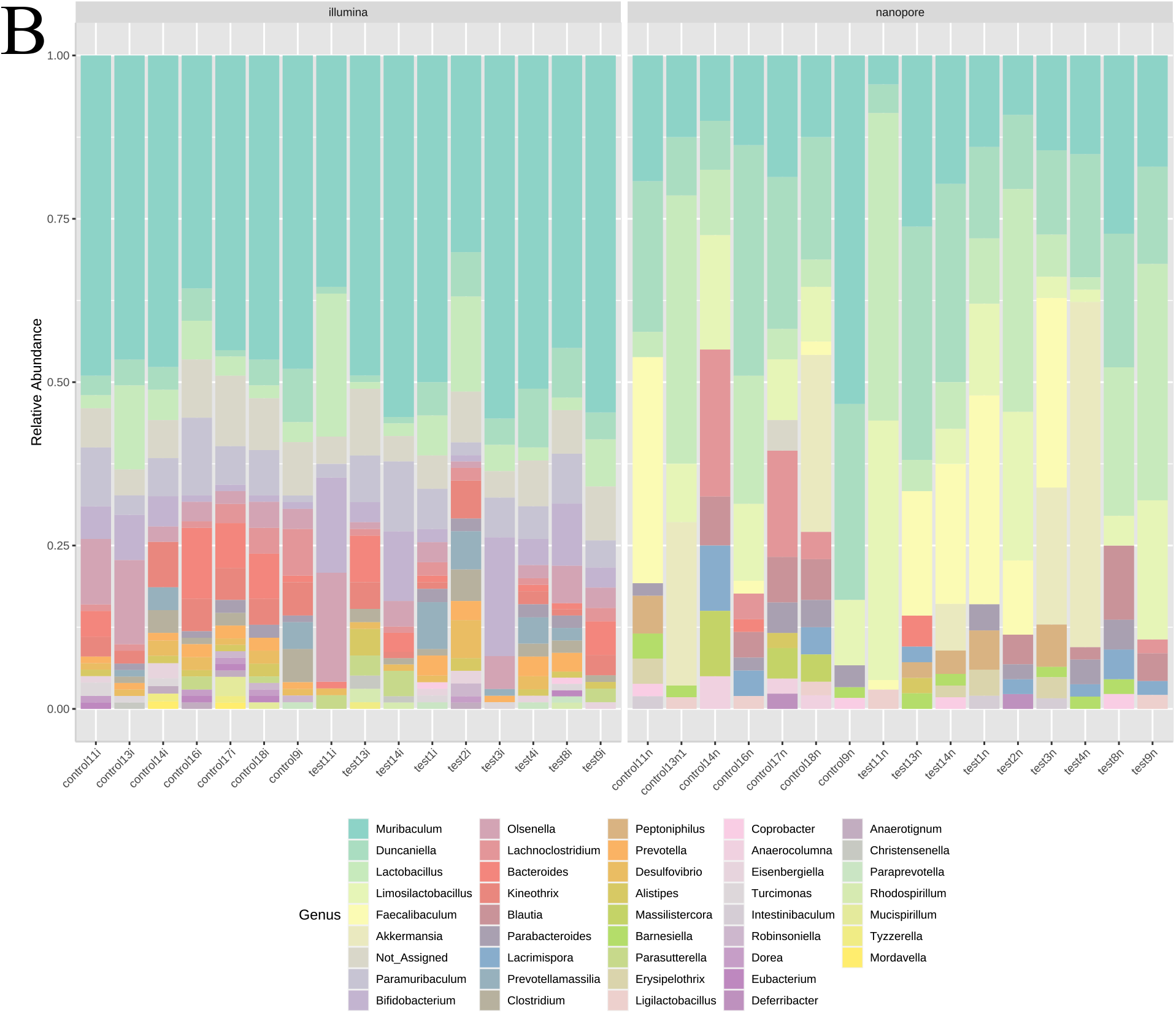
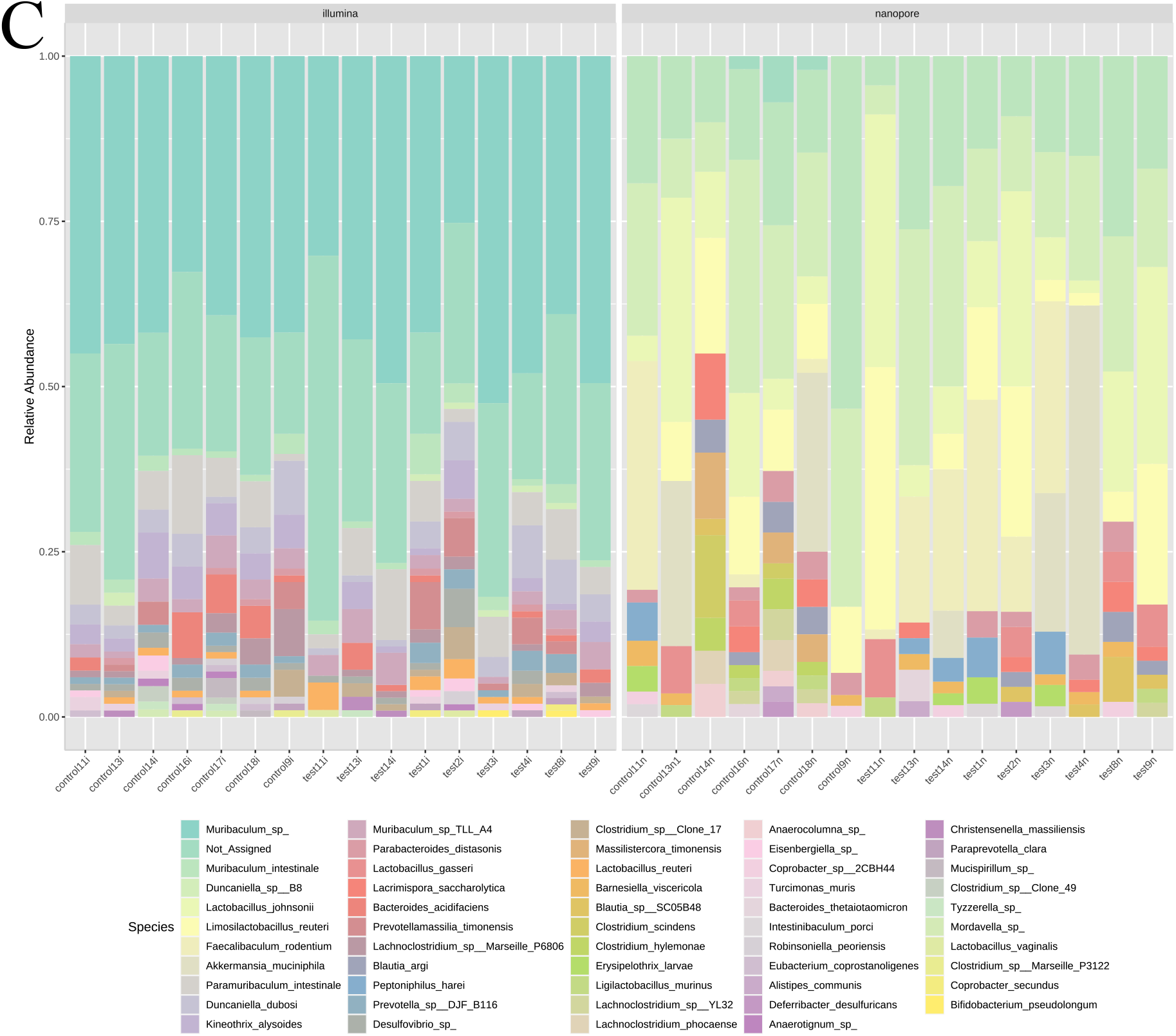
Stacked bar plot displaying percentage abundances comparison between nanopore and Illumina sequencing approaches of assigned taxa at the phylum (A), genus (B), and species level (C). Right: log-transformed relative abundances, and species level (C).

When analysing the infant gut microbiota by Illumina sequencing, in the ethanol-exposed group *Bacteroidetes* was the most abundant phylum represented (70%) followed by *Firmicutes* (14%), *Actinobacteria* (12%), and *Proteobacteria* (3%). *Bacillota* and *Deferribacteres* did not exceed 1% relative abundance and were only observed in three and four samples, respectively. Among the data, 0.5% of the detected reads was not assigned to any phylum. At the genus level, *Muribaculum* accounted for most of the reads observed (47%), followed by *Bifidobacterium* (7%), *Lactobacillus* (7%), *Paramuribaculum* (6%), *Olsenella* (5%), and *Duncaniella* (4%). Around 6% of the reads did not have any genus attributed. The remaining 37 genera were represented by less than 3% of the reads each. Amongst all reads, 39 species were observed and 26% of the reads were not resolved to the species level. The top species detected were *Muribaculum* sp. (42%), *Paramuribaculum intestinale* (6%), and *Duncaniella dubosi* (4%). The remaining 36 species observed represented less than 3% each.

In the control group *Bacteroidetes* was the most abundant phylum represented (68%), followed by *Firmicutes* (18%), *Actinobacteria* (8%), and *Proteobacteria* (3%). *Deferribacteres* and *Bacillota* did not exceed 1% relative abundance and were only observed in five and one sample, respectively. Among the data, 0.6% of the detected reads was not assigned to any phylum. At the genus level, *Muribaculum* accounted for most of the reads observed (46%), followed by *Paramuribaculum* (6%), *Olsenella* (5%), *Lactobacillus* (5%), *Kineothrix* (4%), *Bacteroides* (4%), and *Duncaniella* (4%). Around 7% of the reads did not have any genus attributed. The remaining 33 genera were represented by less than 3% of the reads each. Amongst all reads, 42 species were observed and 23% of the reads were not resolved to the species level. The top species detected were *Muribaculum* sp. (41%), *P. intestinale* (6%), *Kineothrix alysoides* (4%), and *D. dubosi* (4%). The remaining 37 species observed represented less than 3% each.

Illumina sequencing detected a higher number of reads assigned to *Bacteroidetes* in comparison to nanopore sequencing (70% *vs* 51%), and less reads assigned to *Firmicutes* (16% *vs* 50%). Additionally, no reads were assigned to *Verrucomicrobia* (comparted to 9% detected by nanopore sequencing), and 10% of the reads were assigned to *Actinobacteria* where in the nanopore sequencing approach this phylum was not detected. An average of 24% (control group) and 27% (ethanol-exposed group) of the reads were not assigned to the species level in Illumina sequencing. Both ethanol-exposed and control groups showed similar profiles at the phylum, genus, and species level. Nonetheless, there were detected some minor differences in taxa exclusively found in each of the experimental groups. However, those taxa were only present in a limited number of samples and in a very low abundance. In the ethanol-exposed group, the exclusive taxa were *Rhizobium* sp. UB12 (n = 2, relative abundance < 1%), *Gemella* sp. (n = 2, relative abundance < 1%), and *B. pseudolongum* (n = 2, relative abundance < 1%); and in the control group, *Mordavella* sp. (n = 3, relative abundance < 1%)., *M. massiliensis* (n = 4, relative abundance < 1%), *Marvinbryantia* sp. (n = 2, relative abundance < 1%), *Clostridium* sp. C5 48 (n = 2, relative abundance < 1%), *Clostridium* sp. Clone 49 (n = 3, relative abundance < 2%), and *Clostridium* sp. Marseille P3122 (n = 4, relative abundance < 1%).

Only three species were detected in both sequencing platforms, *M. intestinale, P. distasonis*, and *D. muris*. Additionally, *M. intestinale* and *D. muris*, were assigned to more than 50% of the reads in both sequencing approaches irrespectively of the experimental group. Therefore, these species may be extrapolated as members of the murine gastrointestinal tract (29) (30) (31). A total of 36 species were exclusively detected by nanopore sequencing and 74 taxa by Illumina sequencing (data not shown). When both datasets were analysed together, no correlations were found between any taxa and ethanol-exposure or between the control group. All the sequencing platform-exclusive taxa detected were correlated with their associated sequencing approach (*rho* > 0.6; p-value < 0.05; FDR < 0.05). The exception was *A. muciniphila* that showed a moderate but significant correlation with the nanopore sequencing approach (*rho* = 0.4; p-value < 0.05; FDR < 0.05).

The identical alpha- and beta-diversity found between ethanol-exposed and control groups reveals that ethanol does not pose a strong effect on the microbial diversity composition found in the infant mice guts.

In the control and ethanol-exposed groups two taxa were detected in more than 50% of the samples analysed by the two sequencing platforms: *M. intestinale* and *D. muris*. These two species showed a positive correlation with the nanopore sequencing approach (*rho* > 0.6: p-value < 0.05; FDR < 0.05), which may be considered a sequencing bias (Figure 2).

**Figure 2.**
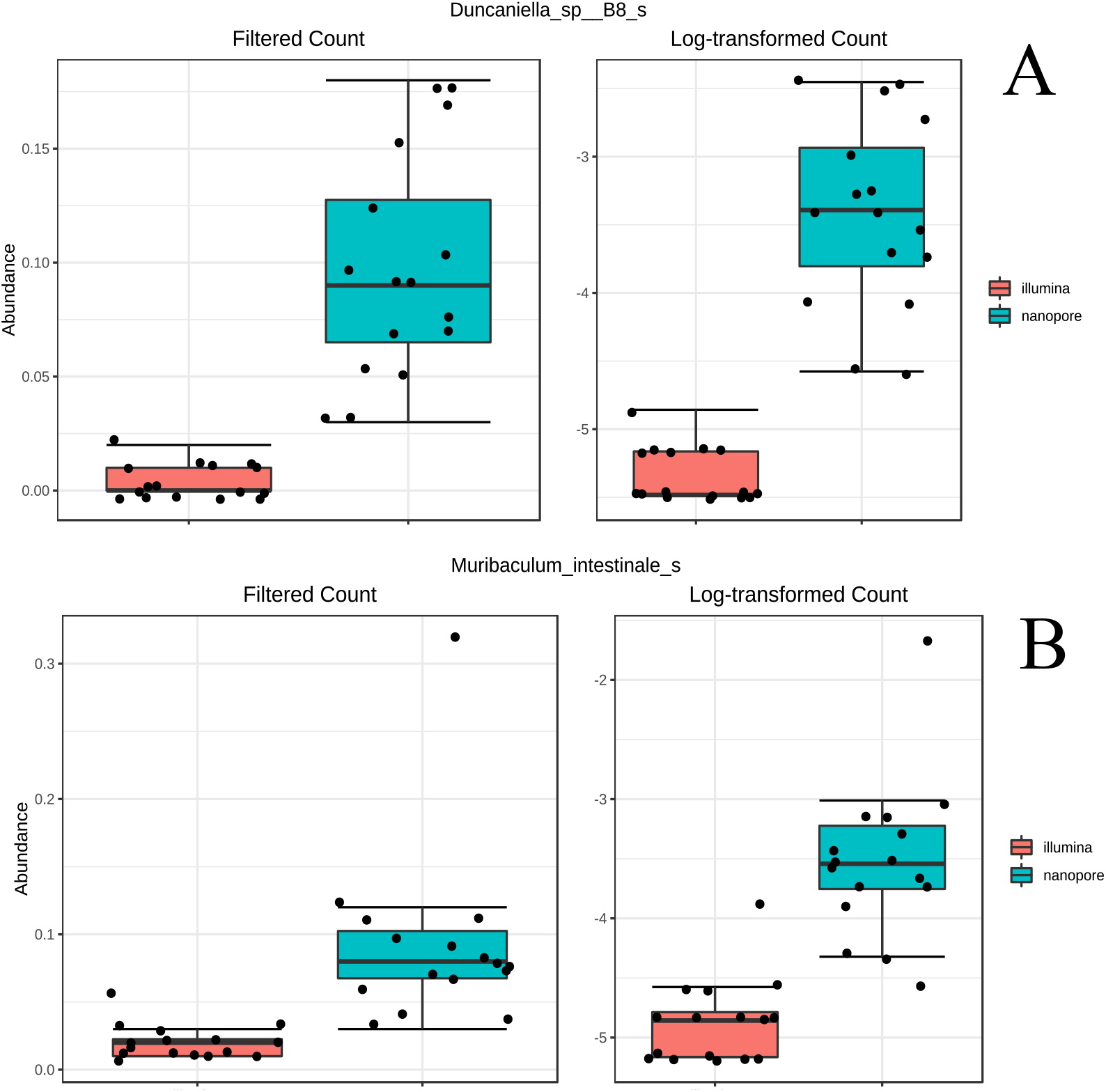
Boxplot displaying the relative abundances of *D. muris* (A) and *M. intestinale* (B) (Spearman correlation coefficient > 0.8; p-value < 0.05; FDR < 0.05) in the two sequencing approaches. Left: raw relative abundances; Right: log-transformed relative abundances.

There were differences detected in the abundance profiles of several taxa at the phylum, genus, and species level when comparing both sequencing platforms (integer Log LDA score > 2; p-value < 0.05: FDR < 0.05) (Figure 3). Due to the overrepresentation of *Bacteroidetes* phylum in Illumina sequencing, this taxon may work as a marker (sequencing bias) when Illumina sequencing is performed. This observation can be explained by high number of reads assigned to *Muribaculum* spp.

**Figure 3.**
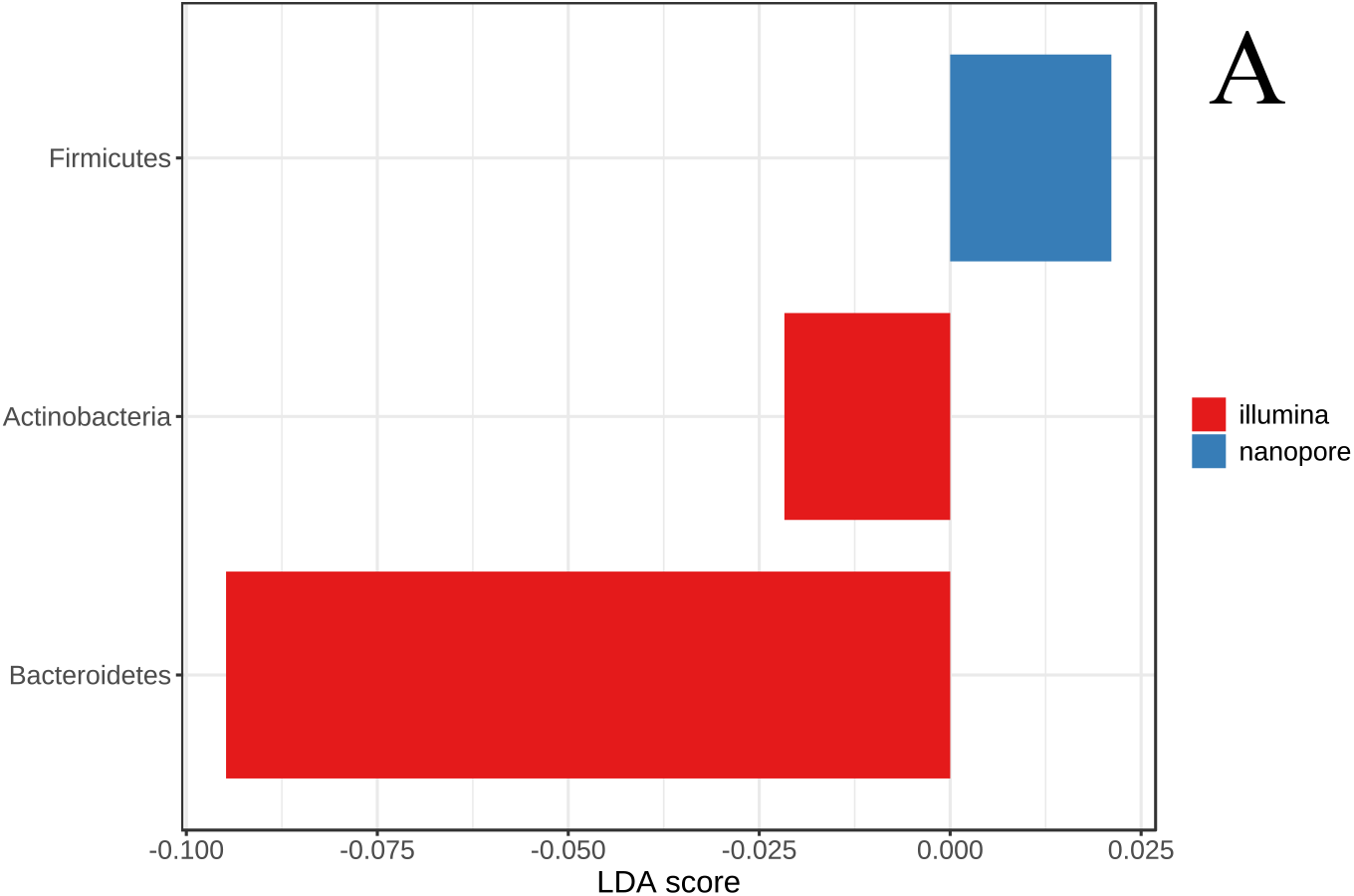
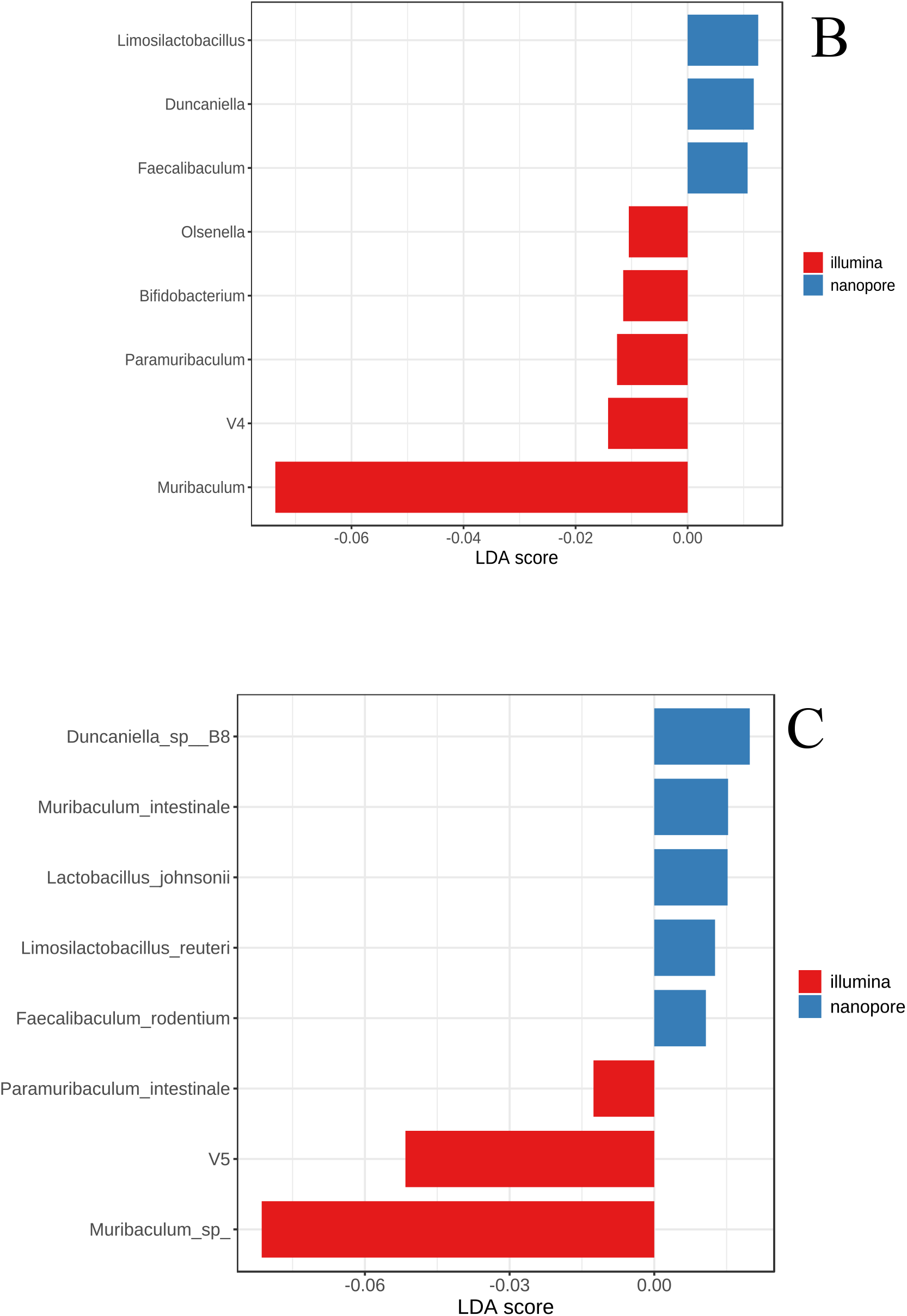
Linear Discriminant Analysis (LDA) Effect Size (LEfSe) showing the relevant taxa significantly abundant and associated with nanopore or Illumina sequencing approaches. The length of each bar corresponds to the LDA score cutoff of 2 (A, phylum level), 1 (B, genus and C, species level); (p-value < 0.05). ‘V4’ and ‘V5’ correspond to the relative abundance attributed to the non-assigned reads at the mentioned taxonomical rank level.

## Discussion

The absent fungal PCR amplification here reported is unexpected. One explanation might be the suboptimal PCR composition and cycle conditions which were optimised for a set of fungal species that may not be found in the mouse gut samples (32). However, the hypothesis of the fully absent mice gut mycobiome cannot be discarded, due to the highly transient feature of the gut fungi members in the mouse model under investigation in this study (33). In this work, infant mice were kept under a controlled environment, which perhaps did not give chance to development of any gut mycobiome at all at early age.

Not many reports have explored nanopore sequencing to microbial profile mice gut microbiomes. One of such studies were performed by Shin and colleagues (2016), who evaluated the performance of a combined approach of nanopore and Illumina sequencing. The results here showed significant differences with the microbial profile obtained by Shin and colleagues (34). Only genera *Lactobacillus* and *Akkermansia* were matched between this study theirs. Since Shin and colleagues’ report dates to 2016, advances in the sequencing chemistry embedded in the flowcell apparatus (FLO-MAP103 *vs* current FLO-R9.4.1 now employed) and the overall improvement of the nanopore sequencing-related bioinformatics pipelines (Metrichor *vs* current *guppy* basecallers) can probably explain the observed differences.

One species showed different relative abundance between the experimental groups in the nanopore sequencing experiment. In accordance with other reports, *F. rodentium* was detected in 13% of the reads in the ethanol-exposed group, and in the control group about 5% of the reads were assigned to the same taxon (4). The clinical relevance of this taxon is being increasingly studied due to its importance in triggering mice depression (35), protection of intestinal tumour growth (36), and further associations within the gut-bran axis (35).

Additionally the microbial profiles detected by Illumina sequencing do not match the previous reports and the disparities may be explained by the different experimental conditions here performed (11) (27). Specifically, neither *Pediococcus, Faecalibacterium, Escherichia*, nor *Staphylococcus* were detected in this study, which was also reported by others (25). In the microbial profile retrieved by the Illumina sequencing, the only genera found that matched the ones detected by Shin and colleagues (2016) were *Lactobacillus* and *Bacteroides*. One of the major differences was the detection of genera *Muribaculum* and *Paramuribaculum*, which were abundantly detected in this study and absent from their report (34).

Amongst the low prevalent taxa detected by Illumina sequencing, the presence of *Rhizobium* sp. may constitute a contamination or a misassignment as previously observed with other taxa, since it was observed with relatively low abundance. *Gemella* spp. have been previously collected from the human microbiome and potential representing pathogenic risk (37) (38) (39). *Mordavella* sp. is a species that was previously isolated from a faecal sample obtained from a healthy volunteer (40) and its presence in laboratory mice gut samples has been linked to food contamination (41). *M. massiliensis* is a species that have been detected in the human colon right side, however, its characterization is still insufficient to draw any conclusion about its importance to the human or mouse host (42). Recently *M. massiliensis* was associated with a dysbiotic state in an obesity mouse model (43). *Marvinbryantia* spp. have been associated with brain lesions in mice, increasing in abundance after traumatic brain injury (44). In humans, these taxa were observed with decreased abundance in faecal samples collected from hypertension patients and showed to be correlated with intestinal vitamin D, which is a risk factor for hypertension (45).Similarly to what was observed with nanopore sequencing, some taxa were exclusively detected in one of the experimental groups by illumina sequencing, although showing relatively low abundance. The exceptions were the genus *Bifidobacterium* that were only abundant in the control group. Contrary to some previous independent reports performed in humans, the control group showed a higher prevalence of *Bifidobacterium* (9) (46) (47). However, other reports observed similar *Bifidobacterium* abundances in the control/healthy subjects as here reported (3) (48) (49) (50) (51). Since *Bifidobacterium* has a known potential to accumulate acetaldehyde under ethanol presence (9) by encoding an acetaldehyde dehydrogenase (51), the results presented in this study reveal that this taxon may have a more complex phenotypic regulation under ethanol presence than previous thought (9) (46) (52).

Besides *D. muris, M. intestinale*, and *P. distasonis*, the remaining taxa exclusively detected by each sequencing approach may be considered members of the rare gut microbiota due to their low relative abundance (53) (54) (55) (56) (57) (58) (59). However, the hypothesis of erroneous taxonomical assignment of those sequence reads cannot be ruled out, specifically in the nanopore sequencing approach. The differences detected in the reads assigned to *Muribaculum* spp. are most probably explained by the distinctive sequencing experimental and/or bioinformatic procedures.

Interestingly, only the nanopore sequencing detected *A. muciniphila*. This observation is contrary with the observation of Shin and colleagues (2016) where they detected *A. muciniphila* using Illumina and nanopore sequencing. Nevertheless, *A. muciniphila* prevalence differed in the ethanol-exposed group (4-28%), which might implicate that ethanol imposes stress on this species. Thus, *A. muciniphila* may have a relevant role in ethanol addiction. This hypothesis is supported by other reports that stated that increasing ethanol habits decrease the abundance of *A. muciniphila* in humans (60) (61). Additionally, the functional capacity of this species to produce/metabolize ethanol (62), and its role in the inflammation of the gut barrier (63) makes the abovementioned hypothesis essential to be investigated. However, caution must be made when using mice models to study the host-microbe interactions of *A. muciniphila*, because despite being considered a commensal member of the human gastrointestinal tract, recent reports have supported the hypothesis that *A. muciniphila* beneficial properties and specific responses are context dependent and differ between human and mice (64). Still, the nanopore sequencing platform basecalling error-rate should be taken into consideration because it may under or overrepresent the true prevalence of this and other affiliated taxa. These observations constitute a major difference between the nanopore and Illumina approaches and, thus, the capacity of each platform to truly assign the correct taxonomy of reads obtained from certain taxa, such as *A. muciniphila*.

Illumina sequencing assigned the majority of the reads to *Muribaculum* sp., more than twice of the reads that nanopore sequencing approach assigned to *M. intestinale* alone (42% *vs* 18%). The distinctive experimental and bioinformatic pipelines performed by Illumina and nanopore sequencing platforms can cause major discrepancies in terms of the microbial profile obtained while analysing the same samples. Here, Illumina sequencing overrepresented *Muribaculum* sp. while nanopore sequencing retrieved less than half of that number of reads to *M. intestinale*. Additionally, nanopore sequencing attributed much more reads to *M. intestinale* than Illumina sequencing (18% *vs* 4%). In the *Muribaculaceae* family it was observed disparities that reveal the taxonomical discrimination capacity of either sequencing platforms. Indeed, nanopore sequencing was able to discriminate more species than Illumina sequencing, which concentrated the assignments as *Muribaculum* sp.. This observation supports the hypothesis that nanopore sequencing is more discriminatory than Illumina sequencing in assigning taxonomy identifiers to lower taxonomical ranks (24).

*M. intestinale, P. distasonis*, and *D. muris* are perhaps members of the infant mice core gut microbiota. However, *M. intestinale* and *D. muris* were positively correlated with the nanopore sequencing approach. Therefore, further investigation must be performed to study the relevance of these taxa to the host physiology. Since both species are members of the C57BL/6J mice gut (65) (66) their increased presence in the ethanol-exposed group may be interpreted as a response to the ethanol stress. The *M. intestinale* role in the mouse gut physiology is not totally understood, however, this anaerobic species can degrade galactose (67) and it is associated with homoserine and serine metabolism (68), thus showing potential to impact the mouse gut homeostasis (69). *D. muris* have been negatively correlated with fat-diet in mice (70), and it was reported as a protection effector against colitis and gut epithelium injury in a mouse model of dextran sulphate sodium-induced injury (71).

*Prevotella* and *Bifidobacterium* are commonly found in the mice gut microbiota (25). Thus, the incapability of nanopore sequencing to detect these taxa represent a substantial handicap. Several taxa were detected with low relative abundance, and they may be considered as members of the rare gut biosphere. The relevance of the rare gut biosphere for the host homeostasis must be studied taking into account every species’ individual and combined effect (53) (54) (55) (56), Lim 2018, de Gouffau 2019). Commonly, these rara taxa represent the majority of species found in environmental studies, displaying unique ecology and biogeography (54). Additionally, the rare biosphere represents the broad diversity and encompasses a wide ecological function arsenal and resiliency, providing the habitat a functional redundancy and flexibility (54) (56). However, sequencing noise and biological artifacts may also contribute to the incorrect taxonomical assignment of part of the rare biosphere composition (54).

In conclusion, there were several taxa correlated with the sequencing approaches, but such correlations may lack biological meaning due to their exclusiveness detection in one the sequencing approaches. A high number of taxa were exclusively obtained by each sequencing platform. However, these sequencing platform-exclusive taxa were not abundantly observed. Therefore, these taxa may be considered rare taxa, and their presence can potentially be explained due to either lab-related or bioinformatic-related experimental errors or limitations (54) (55) (57) (58) (59). Despite the differences found in the microbial profiles and relative abundances of both experimental groups by Illumina and nanopore sequencing, there were no significant differences detected in the diversity indices within samples. The beta-diversity differences were more robust at lower taxonomical ranks, which mean that the microbial profiles obtained by Illumina or nanopore sequencing approaches greatly differ. Despite the differences between sequencing methodologies, *M. intestinale*, and *D. muris* were detected by both approaches, therefore validating their presence in the mice gut since early age. Both sequencing platforms validated the same conclusion in terms of the effect of the ethanol-exposure *in utero*. Ethanol exposure did not affect the microbial profiles. Nevertheless, further studies must be performed to detail the effect of ethanol exposure to target taxa like *A. muciniphila*, since no clear pattern was detected throughout this study.

## Materials and Methods

### Ethical Statement

The mice experimental design was licenced under the Animals (Scientific Procedures) Act, 1986), with license number PPL P5686A14D. All experimental work was performed in accordance with the ARRIVE guidelines published by the National Centre for the Replacement Refinement & Reduction of Animals in Research. Animal Welfare and Review Board ethically approved all protocols and procedures.

### Animal husbandry

Three male and three female C57BL/6J mice (six and eight weeks old, respectively) were randomly paired into three groups. Each male randomly paired with one female and placed into an Optimice® cage. Three monogamous breeding pairs were achieved: BOX1 ethanol (n = 2), BOX2 ethanol (n = 2) and BOX3 ethanol (n = 2). A first batch of 9 infant mice exposed to ethanol *in utero* and 7 non-exposed mice were used for the nanopore and Illumina sequencing. Cage and holding room environment were maintained constantly with controlled light intensity, relative humidity, room and cage temperature. To promote crepuscular behaviour, a 24 h lighting cycle of 12 h light followed by 12 h dark photoperiod was implemented in order to regulate both circadian rhythm and reproductive cycles. Enviro-Dry® fresh nesting was aided and preserved by microenvironment temperature. RM No.3 Special Diet Services (5 g/day/mouse) was supplied to each group.

### Drug formulation and administration

Formulation of 5% ethanol solutions (1000 mL) were formed prior to drug administration and stored at 4 °C. To mimic average human ethanol consumption, for each group, 5% ethanol solution was orally administered from colony set-up onwards, by means of Optimice® internal bottle assembly and allowed fresh daily consumption. Offspring pups were weaned 23-28 days postnatal, separated and placed into gender specific boxes categorised with date of birth, wean date, and box origin.

### Infant mice gut DNA extraction and amplicons generation Fungal gut microbiota

For DNA extraction, MasterPureTM Yeast DNA Purification Kit, according to the manufacturer’s protocol (Lucigen). Extracted DNA was stored at −20 °C until further use. The PCR mixture and thermal profile performed was described above. PCR products were visualized in an agarose gel 1.1% (w/v). Polymerase Chain Reaction (PCR) was performed to amplify the ITS region with diluted genomic DNA (1:10) from each isolate, with the following primer sequences: ITSF1 – CTTGGTCATTTAGAGGAAGTAA; ITS4 – TCCTCCGCTTATTGATATGC. The PCR mixture (25 μL total volume) contained between 1 and 4 ng of DNA template (or 6 μL of unquantified initial DNA), 1× Standard Taq Reaction Buffer, 0.2 mM of dNTPs, 0.4 μM of ITS1F and ITS4, and 0.625 U of Taq DNA Polymerase (New England Biolabs). The PCR thermal profile consisted of an initial denaturation of 1 min at 95 °C, followed by 35 cycles of 30 s at 95 °C, 40 s at 50 °C, 60 s at 68 °C, and a final step of 5 min at 68 °C. The PCR products were visualized in an agarose gel 1.1% (w/v).

### Bacterial gut microbiota

DNA extraction was performed using around 300 mg of biological faecal material with QIAmp PowerFecal Pro DNA Kit, according to the manufacturer’s protocol (Qiagen). The final supernatant containing the extracted DNA was used immediately or stored at −20 °C until further use. DNA quantity and quality (A260/280 and A260/230) were measured using a Qubit 4 Fluorometer (Thermofisher), and with DeNovix DS-11 Spectrophotometer (DeNovix), respectively. PCR was performed to amplify the full-length of the 16S rRNA gene (~1,500 bp). The primers pairs used were the 16S-27F – AGAGTTTGATCCTGGCTCAG, and the 16S-1492R – GGTTACCTTGTTACGACTT.

The PCR mixture for the full-length 16S rRNA gene (25 μL total volume) contained between 0.2 and 1.6 ng of DNA template (or 3 μL of unquantified initial DNA extract), 1× Standard Taq Reaction Buffer, 0.5 mM of dNTPs, 0.4 μM of 16S-27F and 0.8 μM 16S-1492R, and 0.625 U of Taq DNA Polymerase (New England Biolabs). The PCR thermal profile consisted of an initial denaturation of 30 s at 95 °C, followed by 30 cycles of 20 s at 95 °C, 20 s at 51 °C, 2 min at 68 °C, and a final step of 5 min at 72 °C. The amplicons were purified with the AMPure XP beads (Beckman Coulter) using a 0.5× according to manufacturer’s protocol. The quantity and quality of the purified DNA was measured by Qubit 4 Fluorometer, and with DeNovix DS-11 Spectrophotometer, respectively. If the PCR purified amplicons did not show A260/280 between 1.8-2.0 and A260/230 between 2.0-2.2, they were discarded, and the PCR reaction repeated.

### MinION sequencing, basecalling. and bioinformatics analysis

Nanopore sequencing was performed using MinION MK1B sequencer (ONT) and flowcells R9.4.1 (FLOW-MIN106D, ONT) using MinKNOW™ software v.21.02.1 (ONT). After the MinION Flow Cell was setup accordingly to manufacturer’s instructions, the pre-sequencing mix was prepared accordingly to manufacturer’s protocol. Once the DNA sequencing library was loaded, the 24 h sequencing run was initiated. For the targeted sequencing approach using the full-length 16S rRNA gene, four MinION Flow Cells were used to maximize the sequencing output. All nanopore sequencing experiments ran for 24 consecutive hours. Basecalling was performed with *guppy* standalone tool (https://nanoporetech.com) from the command line using the *fast basecalling* mode, with the quality score filter of 8, and adapters’ trimming mode *on*. Quality controls were performed using *nanoplot* tool (72) with the length filters parameters of maximum length at 2000 and minimum at 1000. WIMP (version 2021.11.26) was used for the classification assignments steps.

### Microbial profiling of infant mice gut samples exposed to ethanol *in utero* through Illumina sequencing

Seven control and nine test gut samples from mice infants exposed to ethanol in utero were used for Illumina sequencing (the same samples were formerly analysed with nanopore sequencing). Genomic DNA was extracted from infant mice faecal samples and sent to Eurofins (Germany) for sequencing by Illumina MiSeq platform (Illumina) targeting the V3-V4 region of the 16S rRNA gene. Then, reads were demultiplexed and primers clipped before reads were merged. All reads undergone a quality filter step before microbial profiling analyses. FASTQ files with quality filtered reads were used as input for microbial profiling. Chimeric reads were identified and removed based on the de novo algorithm of UCHIME (73) as implemented in the VSEARCH package (74). The remaining set of high-quality reads was processed using minimum entropy decomposition (75) (76). Minimum Entropy Decomposition provides a computationally efficient means to partition marker gene datasets into OTUs. To assign taxonomic information to each OTU, DC-MEGABLAST alignments of cluster representative sequences to the sequence database were performed. The most specific taxonomic assignment for each OTU was then transferred from the set of best-matching reference sequences (lowest common taxonomic unit of all best hits). Sequences identities of 70% across at least 80% of the representative sequence was set as a minimal requirement for considering references sequences (reference database: /dbdir/nt. ltered.fa; Release 2020-02-03). Further processing of OTUs and taxonomic assignments was performed using QIIME software package (version 1.9.1). Abundances of bacterial taxonomic units were normalized using lineage-specific copy numbers of the relevant marker genes to improve estimates (77).

### Statistical analysis

For comparative analyses, non-normalized relative frequencies were used as input for MicrobiomeAnalyst software (78), and only taxa with relative abundances above 0.1% were considered.

### Diversity index analyses

Alpha-diversity measure was employed to analyse the diversity within a sample or an ecosystem using Shannon-Wiener and/or Simpson indexes. Beta-diversity was employed to measure the diversity between samples using non-metric multidimensional scaling (NMDS) as ordination technique. For community analyses, Bray-Curtis dissimilarity statistic were performed as distance methods.

### Testing for significant differences between groups

After representing objects in ordination plots and assess eventual clustering of similar objects, a statistical test was performed to assess differences between similarities within and between groups. Permutational analysis of multivariate dispersions (PERMDISP), Permutational analysis of variance (PERMANOVA), and Analysis of similarities (ANOSIM) were employed as stated in the results section.

### Clustering and correlation

Cluster analyses were performed in MicrobiomeAnalyst software (78) using Bray-Curtis dissimilarity distance measure and Ward as the clustering algorithm. Although both Spearman and Pearson correlation coefficients might be employed to microbiome data – due to independence of variables under test – the non-parametric Spearman correlation coefficient (*rho*) was employed with an alpha-level of 0.05.

### Comparison and classification

Non-parametric Linear Discriminant analysis (LDA) effect size (LEfSe) was employed for biomarker discovery and data explanation in the Illumina and nanopore sequencing microbial profiling approaches (79) incorporating the non-parametric Kruskal-Wallis test with an alpha-value of 0.05.

## Acknowledgements

This research was supported by the DTA3/COFUND PhD Fellowship Programme (funding from the European Union’s Horizon 2020 research and innovation programme under the Marie Skłodowska-Curie grant agreement No. 801604). The funder had no role in study design, data collection and interpretation, or the decision to submit the work for publication.

## Author contribution

Conceptualization, C.P-R., F.G., L.B., and J.I.; Methodology C.P-R, F.G., L.B., and J.I.; Investigation, C.P-R.; Mice experimental work, N.B.; Supervision, F.G., L.B., and J.I.; Writing – original draft, C.P-R.; Writing – review & editing, F.G., N.B., L.B., and J.I.

## Notes

### Competing Interest Statement

The authors have declared no competing interest.

